# Convergent evolution of sperm gigantism and the developmental origins of sperm size variability in *Caenorhabditis* nematodes

**DOI:** 10.1101/050252

**Authors:** Anne Vielle, Nicolas Callemeyn-Torre, Clotilde Gimond, Nausicaa Poullet, Jeremy C. Gray, Asher D. Cutter, Christian Braendle

## Abstract

Sperm cells provide crucial, if usually diminutive, ingredients to successful sexual reproduction as the source of centrioles and half the diploid genome. Despite this essential conserved function, sperm competition and coevolution with female traits can drive spectacular change in size and shape of these motile cells. Here we characterize four repeated instances of convergent evolution of sperm gigantism in *Caenorhabditis* nematodes using phylogenetic comparative methods on 26 species. Species at the extreme end of the 50-fold range of sperm-cell volumes across the genus have sperm capable of comprising up to 5% of egg-cell volume, representing severe attenuation of the magnitude of anisogamy. Exploring potential genetic and developmental determinants of *Caenorhabditis* sperm size variation, we uncover significant differences in mean and variance of sperm size among genotypes, between sexes of androdioecious species, as well as within and between individuals of identical genotypes. We demonstrate that the developmental basis of sperm size variation, both within and between species, becomes established during an early stage of sperm development, i.e. at the formation of primary spermatocytes while subsequent meiotic divisions contribute little further sperm size variability. These findings provide first insights into the developmental determinants of inter-and intraspecific sperm size differences in *Caenorhabditis*. Together, our results provide a novel integrative view on the developmental and evolutionary origins of *Caenorhabditis* sperm size variation. We hypothesize that life history and/or ecological differences among species favoured the evolution of alternative sperm competition strategies toward either many smaller sperm or fewer larger sperm, with gigantic sperm potentially providing a means of paternal care via gametic provisioning or as a potent vehicle for sexual conflict over offspring development.

## INTRODUCTION

Post-copulatory sexual selection that manifests as sperm competition and cryptic female choice can drive rapid trait evolution in gametes and genitalia. Rapid evolution of such reproductive traits [1, 2] is a hallmark of species differences and can accelerate the accumulation of reproductive isolation in species diversification [3, 4]. Cases of exaggerated phenotypic evolution in sperm size, as for sperm gigantism in some *Drosophila* and other organisms [5-8], provide some of the most dramatic examples in the evolution of form and function. Moreover, exaggerated male gamete formation as a consequence of sperm competition marks an evolutionary reversal in relative male investment away from extreme anisogamy. One perspective on this outcome holds that such increased male investment in offspring represents a means of paternal care via gametic provisioning [8]. Alternatively, incorporation of more male gamete cytoplasm into the zygote might set the stage for greater parental sexual conflict over offspring development [9, 10]. Sperm evolution thus integrates the fundamental evolutionary processes that define inter-male competition, inter-sexual conflict, and biological diversification all in the context of the exceptionally accessible cell biology and development of a single, discrete cell type.

Nematodes of the genus *Caenorhabditis* have emerged as key model organisms to study both mechanisms and the evolution of sperm competition. In general, nematode spermatozoa usually lack cilia or flagella [11], unlike the sperm cells of insects and mammals, so that changes in sperm size must be coupled to cell volume. In the androdioecious nematode *C. elegans* multiple lines of evidence implicate cell size as a key component of sperm competitive ability: (i) males make larger sperm than hermaphrodites, with male sperm consistently outcompeting hermaphrodite sperm [12, 13], (ii) larger *C. elegans* male sperm are competitively superior and crawl faster [14], and (iii) experimentally enhanced male-male competition leads to the evolution of larger sperm [15, 16]. *C. elegans* males (and hermaphrodites) have small sperm compared to related obligatorily outcrossing species [17, 18], which is thought to reflect part of a ‘selfing syndrome’ in species like *C. elegans* that reproduce primarily by self-fertilization and so experience minimal sperm competition in nature [18, 19]. Inter-species matings also demonstrate the competitive superiority of larger sperm [20-22]. Despite the competitive advantages to males of transferring large sperm to mates when under risk of sperm competition, sperm production rates are slower for *C. elegans* genotypes that make larger sperm [14, 23]. Such trade-offs might limit sperm size evolution, depending on the costs and benefits to sperm size versus number, in combination with mating rates and the probability of paternity in achieving fertilization success.

Despite the extensive theoretical, morphological and experimental literature on sperm competition in sexual selection [24-26], the developmental origins of size variation in sperm traits, such as cell size or flagellum length, remain more elusive. Concerning *Caenorhabditis* nematodes, the genetic control of spermatogenesis is well-characterized in *C. elegans* [27-29], yet the developmental mechanisms underlying sperm size differences, e.g. between hermaphrodites and males, are not understood. During *C. elegans* spermatogenesis, the onset of meiosis involves the formation of primary spermatocytes, characterized by their separation from a syncitial germ cell progenitor pool, during which they increase substantially in cell size [30-32]. Subsequently, the primary spermatocyte divides to form two secondary spermatocytes, which rapidly undergo the second meiotic division resulting in four haploid spermatids and an anucleate residual body [31]. The secondary spermatocytes may either remain connected through cytoplasmic bridges or become separated so that each gives rise to two spermatids and a residual body [31]. Meiotic divisions during *C. elegans* sperm development involve extensive cytokinesis and redistribution of cellular components to residual body and spermatids [31-33]. The rapid meiotic divisions in *C. elegans* go hand in hand with a corresponding cell size reduction from primary spermatocyte to spermatid [31]. Male and hermaphrodite spermatogenesis in *C. elegans* appear equivalent, and it remains unclear how *C. elegans* spermatogenesis generates sperm of distinct size according to sex, genotype, individual, or how evolution has shaped male sperm development of gonochoristic *Caenorhabditis* species that display sperm size divergence [17, 18, 22].

To address these questions surrounding sperm evolution and its developmental basis, we quantified sperm size variation across the *Caenorhabditis* phylogeny covering 26 species [34]. In addition to the well-appreciated convergent evolution of sperm miniaturization in self-fertilizing species, we demonstrate convergent evolution of sperm gigantism in four independent lineages. The presence of gigantic *Caenorhabditis* sperm contributes to a 50-fold range of variation in sperm volume among species, with such sperm capable of comprising 5% of egg cell volume, in contrast to *C. elegans* hermaphrodite sperm that are just 0.2% the size of eggs. We further found substantial sperm size variability within species, between sexes of androdioecious species, as well as between and within individuals of the same genotype. We present experimental evidence implicating primary spermatocyte formation as the key stage establishing the developmental basis of both intra-and interspecific variation in sperm size, with little sperm size variability induced during subsequent cell divisions. Our findings provide first insights into the developmental determinants of sperm size variation and the results of our extended species survey are consistent with the notion that sexual selection has shaped the diversification of *Caenorhabditis* sperm size.

## RESULTS

### Repeated evolution of extremely large male sperm in gonochoristic Caenorhabditis species

We quantified male sperm size across the *Caenorhabditis* phylogeny from measures of spermatid cross-sectional area, demonstrating substantial disparity in sperm size among 26 species that include representatives from all major subgroups in the genus [34, 35] (Figures 1A to 1D). Average sperm size varies >13-fold in cross-sectional area of male spermatids (i.e. >50-fold in spermatid volume), ranging from the tiny 20μm^2^ sperm of the androdioecious *C. tropicalis* to the gigantic 281μm^2^ sperm of the gonochoristic *C. plicata* (Figure 1, Tables S1 and S2). Defining sperm gigantism heuristically as an average sperm size >100 μm^2^, we identified four such species in our survey: *C. plicata, C. drosophilae, C. japonica* and *C. macrosperma*, with no instances of sperm gigantism among the set of species from the *Elegans* group analysed here (Figures 1A and 1B). Applying a multi-peak Ornstein-Uhlenbeck process of phenotypic evolution along the phylogeny [36], we find statistical support for four independent transitions in sperm size in these species toward evolutionary convergence of giant sperm (*c*=4 size shifts to two convergent states, P=0.041; Figures 1B and S1). Our survey of all 26 species is consistent with previous observations that, on average, gonochoristic species make substantially larger sperm than androdioecious species [17, 18]. However, sperm size of several gonochoristic species (*C. nouraguensis, C. yunquensis, C. angaria, C*. sp. 1, *C. castelli*) falls within a similar size range (20-30μm^2^) as the small male sperm of androdioecious species (Figure 1A).

**Figure 1.**
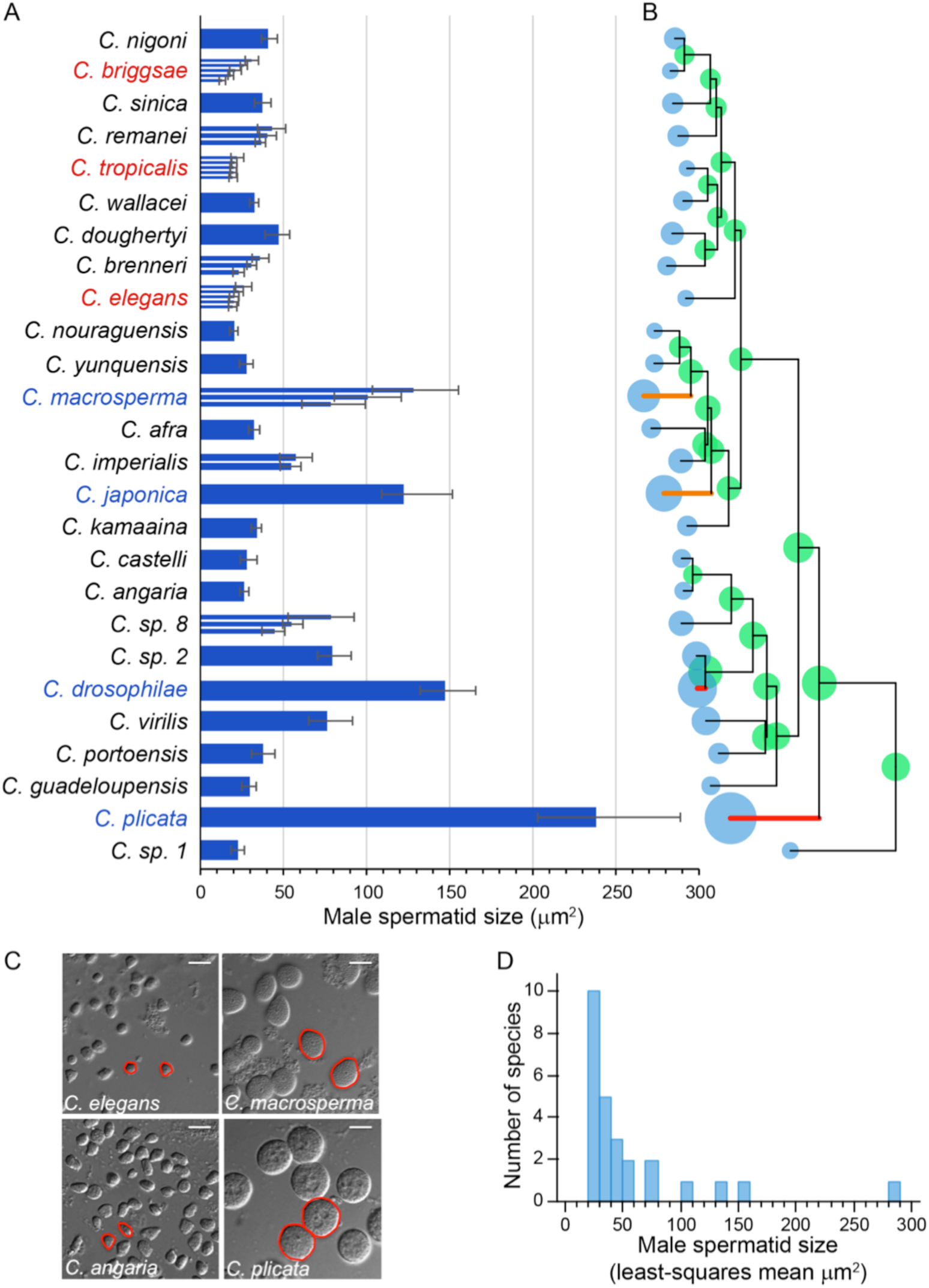
Species differences in *Caenorhabditis* spermatid size show convergent evolution of sperm gigantism. (A) The 26 species show substantial variation in male spermatid cross-sectional area (ANOVA, effect *species*: F_25, 13736_ =3406.40, P<0.0001; per strain: 85-653 spermatids from 4-11 individuals; median ± interquartile range of pooled measurements shown to illustrate range of variation). Narrow bars within a species indicate medians of different wild isolate genotypes. For details of sample sizes and complete statistical results, see Tables S1 and S2. (B) Mapping of extant and ancestral spermatid size on the *Caenorhabditis* phylogeny identifies four evolutionary transitions to gigantic sperm. Lineages highlighted red (*C. plicata*, *C. drosophilae*) and orange (*C. macrosperma*, *C. japonica*) indicate the two convergent sperm size regimes identified by SURFACE under a Hansen model of the Ornstein-Uhlenbeck process of stabilizing selection in trait evolution [36]. The area of circles at each node is proportional to the species least-squares mean spermatid cross-sectional area (blue), with ancestral states (green) inferred from a Brownian motion model of trait evolution in GEIGER [86]. Metrics of phylogenetic signal for sperm size indicate strong dependence of trait values on phylogenetic relationships of the taxa (Pagel’s **A**, = 0.96, Blomberg’s K=0.48). Phylogeny and branch lengths from [34]. (C) DIC images of representative spermatid cells for four species of *Caenorhabditis* illustrate the sperm gigantism of *C plicata* (SB355) and *C macrosperma* (JU1857) relative to *C elegans* (N2) and *C angaria* (PS1010). Scale bars: 15μm. (D) A histogram of sperm size across species (least-squares mean estimates) reveals the outlier trait values for the four species with sperm gigantism.

### Allometry, anisogamy and trade-offs in sperm size evolution

Macro-evolutionary patterns of sperm size disparity could simply reflect a consistent allometric scaling of animal and cell size. However, when we test for coevolution of sperm and male body length (Table S3, Figure S2) using phylogenetic corrections, we find no relationship (log-transformed values; F_1,23_=1.821, P=0.19) (Figure 2A). Measures of egg (embryo) size also differ among species [37], although, in contrast to sperm, the range of variation is less than three-fold (embryo longitudinal area from 859 μm^2^ for *C. angaria* to 2474 μm^2^ for *C. plicata*) (Figures S2 and S3). Sperm size and egg size do correlate positively (Figure 2B), and yet substantial residual variation remains in sperm size among species. Consequently, the evolution of male sperm size yields striking changes in the magnitude of anisogamy among species (Figure 2C), ranging from an egg: sperm volumetric ratio of 453:1 in *C. elegans* to 20:1 in *C. plicata* (Figures 2C to 2E). These findings are consistent with previous conclusions for *Caenorhabditis* that sexual selection by sperm competition is the key driver in the evolution of male sperm size.

**Figure 2.**
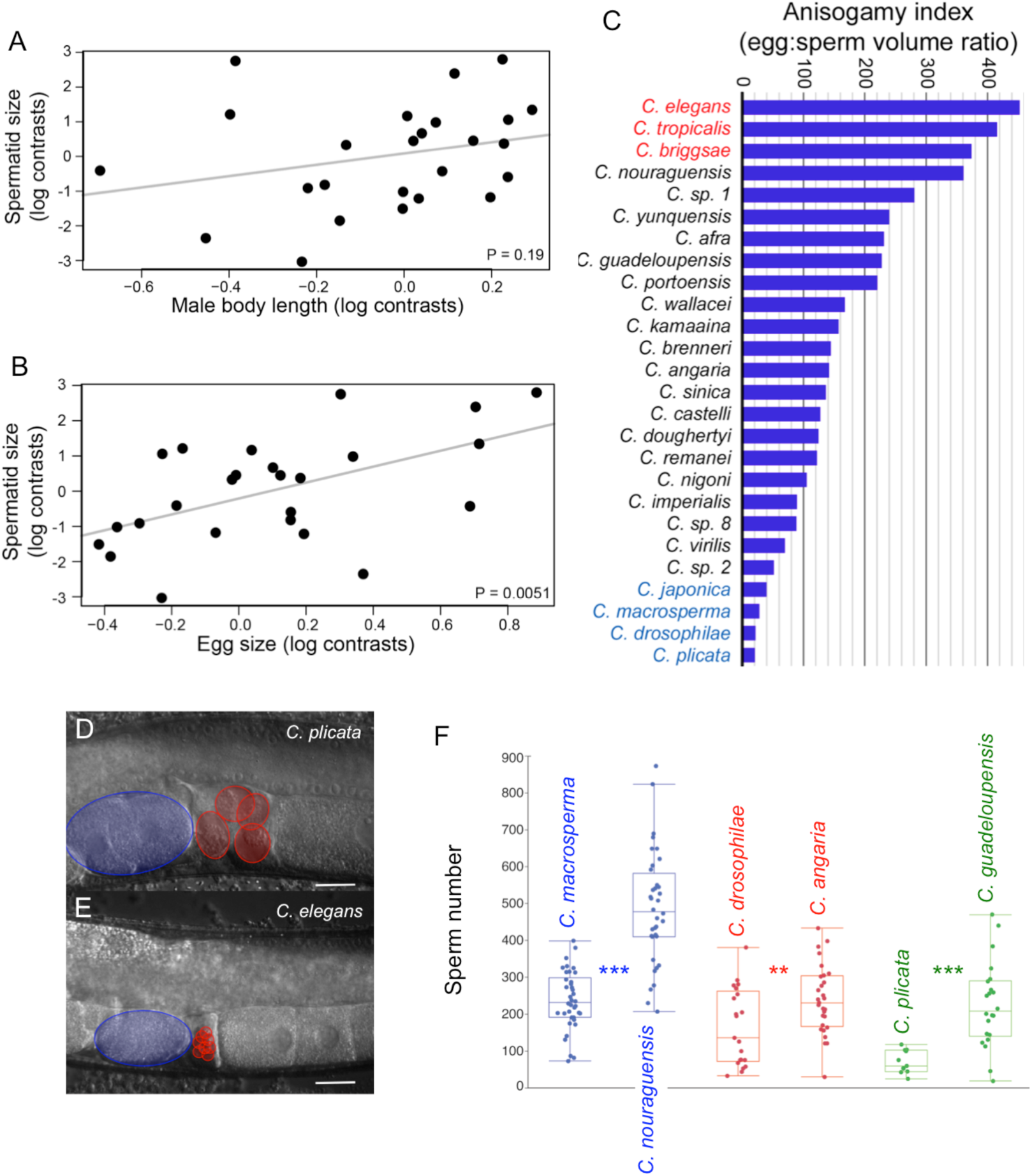
Allometry, anisogamy and trade-offs in sperm size evolution. (A) Species with larger males do not make larger sperm, as indicated in the non-significant slope in this plot of phylogenetic independent contrasts (PIC) of sperm cross-sectional area and male length (log-transformed) (F_1,23_=1821, P=0.19). (B) PIC plot of spermatid size versus egg (embryo) size (log-transformed) shows a significant positive correlation (F_1,23_=9.58, P=0.0051); embryo size data from Farhadifar et al. (2015). (C) Despite variation in egg size and its correlation with male sperm size, an index of anisogamy (embryo volume / spermatid volume) differs drastically across species, with sperm size being responsible for most of the heterogeneity. (D, E) DIC images of the reproductive tract in an inseminated female of *C plicata* (SB355) and a hermaphrodite *C elegans* (N2) with self-sperm, illustrating the extreme species divergence in anisogamy. Spermatozoa in the spermatheca are outlined in red, and an early embryo in the uterus is outlined in blue. Scale bars: 20μm. (F) In three contrasts of species with ‘standard’ versus ‘gigantic’ sperm, the species with larger sperm transfers significantly fewer of them per mating (*C. nouraguensis* vs *C macrosperma*: F_1,72_=82.66, P<0.0001; *C angaria* vs *C drosophilae*: F_1,48_=8.16, P=0.0063; *C guadeloupensis* vs *C plicata*: F_1,31_=21.71, P<0.0001).

Previous work within species of *Caenorhabditis* has demonstrated fertilization advantages to large compared to small sperm [14], even leading to the evolution of larger sperm under experimentally elevated polygamous mating conditions in *C. elegans* [15, 16]. However, if males transferring large sperm are constrained in the number that they can transfer to a female, then such a trade-off could limit the evolution of ever-larger sperm. The evolution of fewer sperm per ejaculate should occur only with greater assurance of paternity, e.g. less polygamy and weaker inter-male sperm competition [38]. To test for a possible fitness trade-off for males having gigantic sperm, we quantified the number of transferred sperm following a single mating for three contrasts of species pairs differing in sperm size. Consistent with the possibility of a size-number trade-off, in each case, the species with larger sperm transferred fewer of them per copulation (Figure 2F). Given the large fraction of the body cavity comprised of gonad tissue, male width also may correspond to investment in testis; for example, increased width of the vas deferens may allow passage of larger sperm and/or increased width of the distal germline may permit growth of larger spermatocytes. Consistent with these possibilities, spermatid diameter can be >25% of the width of male worms in species with sperm gigantism and we indeed observed a significant positive correlation of spermatid size and male body width (phylogenetic contrasts of log-transformed values; F_1,23_=19.21, P=0.00022).

### Developmental origin of sperm size differences among species

Given the enormous disparity in sperm size across *Caenorhabditis* species, what developmental underpinnings at the cellular level might account for the origins of large versus small sperm? In particular, to what extent do initial steps of spermatogenesis, i.e. the formation of primary spermatocytes, diverge to generate sperm size variation? To address this question, we examined spermatogenesis in male gonads of ten species with divergent sperm size (Figure 3). Observation of live animals using Nomarski optics indicates that species producing large spermatids also display large primary spermatocytes (Figure 3A). To obtain quantitative estimates of primary spermatocyte size variation, we measured cell size of male primary spermatocytes in the karyosome stage at the end of meiotic prophase, when chromosomes aggregate into a highly condensed mass [30], using dissected germlines of young adult males stained with DAPI and Phalloidin to visualize nuclei and cell outlines, respectively (Figure 3B). Species variation in primary spermatocyte size variation mirrors spermatid size variation (Figure 3C), and the size distribution of primary spermatocytes strongly predicts spermatid size (phylogenetic contrasts on log-transformed values; F_1,7_=25.65, r^2^_adj_=0.76, P=0.0015) (Figure 3D), indicating that male spermatid size is largely determined by the initial size of primary spermatocytes. We conclude that *Caenorhabditis* sperm size determination occurs during an early stage of sperm development, prior to the detachment of primary spermatocytes from the rachis [30], i.e. prior to meiotic divisions.

**Figure 3.**
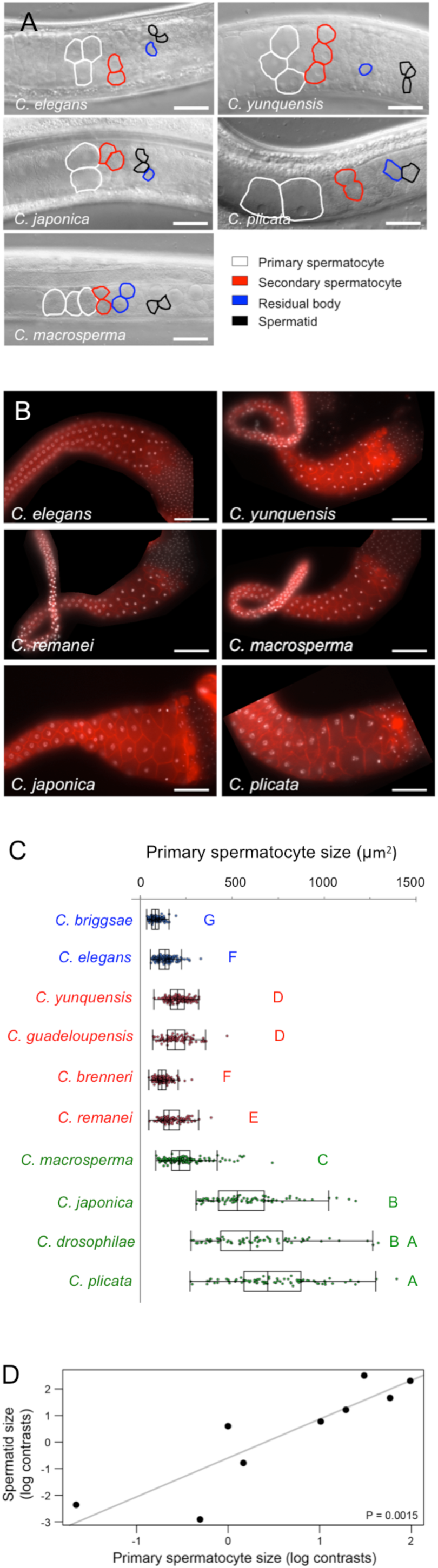
Developmental origin of sperm size differences among species. (A) DIC images of representative male gonads for two species with ‘small’ sperm sizes (*C. elegans, C yunquensis*) and three species with ‘gigantic’ sperm (*C. japonica, C macrosperma, C plicata*), visualizing the transition from primary spermatocytes to spermatids. Scale bars: 20μm. (B) Phalloidin (red) and DAPI (white) stainings of dissected male gonads of select species with divergent sperm size. Scale bars: 40μm. (C) Quantification of male primary spermatocyte size in ten *Caenorhabditis* species with divergent sperm size, arranged by increasing spermatid size from top to bottom. Androdioecious species with the smallest male sperm (blue), gonochoristic species with ‘standard” sperm size (red) and ‘gigantic’ sperm (green). Primary spermatocyte size shows significant variation among species (ANOVA, effect *species*: F_9,1101_=603.88, P<0.0001; for sample sizes and complete statistical results, see Tables S5 and S6). Values labelled with different letters indicate significant differences (Tukey’s HSD, P<0.05). (D) The size of spermatids and primary spermatocytes correlates strongly across examined species (10 species; phylogenetic independent contrast on log-transformed values; r^2^_adj_=0.76, F_1,7_=25.65, P=0.0015).

### Extensive genetic variation for sperm size within species and between the sexes

In addition to assessing sperm size differences among species, we next quantified sperm size variation of multiple wild isolates (genotypes) in gonochoristic and androdioecious species. Measuring male sperm size for isolates of gonochoristic species, we find substantial heritable variation within all four species examined (Figure 4A). For the three androdioecious species, we measured sperm sizes separately for males and hermaphrodites from each of five distinct wild isolates per species, also revealing extensive genetic variation in sperm size for both sexes (Figure 4B). Hermaphrodite sperm are consistently smaller than male sperm in all species including *C. tropicalis* (Figure 4B), consistent with previous studies of *C. elegans* and *C. briggsae* [18, 21]. Hermaphrodite sperm of *C. briggsae* are smallest, and *C. tropicalis* largest, both in absolute size and relative to conspecific males (Figure 4B). We also observed a weak positive correlation between the average sperm size of males and hermaphrodites in *C. tropicalis*, but not in *C. elegans* or *C. briggsae* (Figure 4C). The absence of consistent or strong inter-sexual correlations of sperm sizes suggests that developmental genetic control of sperm size regulation need not be shared between sexes of androdioecious species despite their superficially equivalent spermatogenic developmental programs.

**Figure 4.**
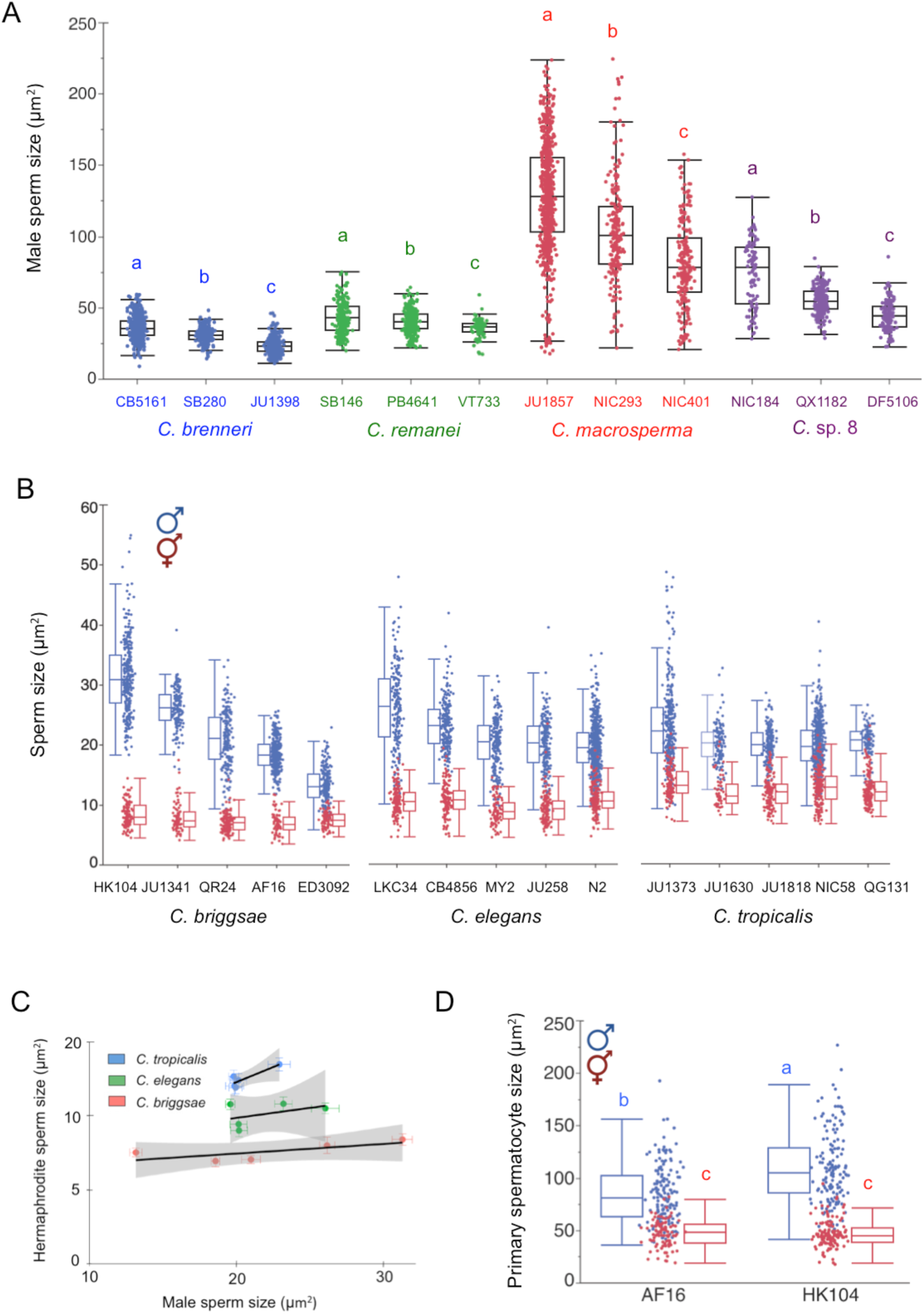
Genetic variation for male sperm size and sperm size differences between males and hermaphrodites. (A) Male sperm size shows extensive genetic variation within each of four gonochoristic *Caenorhabditis* species (ANOVAs performed separately for each species, *C. brenneri*: F_2,1211_=506.88, P<0.0001; *C. remanei*: F_2,599_=27.47, P<0.0001; *C. macrosperma*: F_2,1026_=160.08, P<0.0001; *C*. sp. 8: F_2,450_=147.55, P<0.0001; for complete statistical results, see Table S7). Values labelled with different letters indicate significant differences among isolates within a species (Tukey’s HSD, P<0.05). (B) Distributions of sperm size for hermaphrodites and males of five distinct wild isolates of each of the three androdioecious species. In each species, there is significant genetic variation for both male and hermaphrodite sperm size, and average male sperm size is always greater than hermaphrodite sperm size (for data and complete statistical tests, see Tables S8 and S9). (C) Male and hermaphrodite mean sperm sizes are significantly correlated across isolates for *C. tropicalis* (F_1,3_=15.62, R =0.83, P=0.028) but not *C. elegans* (F_1,3_=1.20, R^2^ =0.29, P=0.35) and *C. briggsae* (F_1,3_=1.42, R^2^ =0.32, P=0.32). (D) Primary spermatocyte size variation of hermaphrodites and males in *C. briggsae* isolates AF16 and HK104. Values labelled with different letters indicate significant differences (Tukey’s HSD, P<0.05; for complete statistical results, see Table S10).

We also found that, in androdioecious species, differences in sperm size between sexes are correlated with corresponding differences in primary spermatocyte size. This observation is indicated clearly in the comparison of *C. briggsae* isolates AF16 and HK104, for which males exhibit significantly larger primary spermatocytes than hermaphrodites (Figure 4D). HK104 males also displayed significantly larger primary spermatocytes compared to AF16 males (Figure 4D), consistent with male sperm size differences between these isolates (Figure 4B). These results indicate that, as for species differences, sperm size differences among genotypes and between sexes can be explained by corresponding size differences in primary spermatocytes.

### Pronounced intra-and inter-individual sperm size variability across species

In addition to the substantial differences in average male sperm size among genotypes and species, we also observed considerable sperm size variability between and within single animals for all isolates examined (e.g. Figures 4A and 4B). Theory predicts that species with stronger sperm competition ought to exhibit lower coefficients of variation (CV; ratio of standard deviation to mean) in sperm traits [6, 39-41]. Therefore we tested for reduced CV in species with larger spermatids, which might be expected if sperm size provides the principal indicator of the intensity of sperm competition in a species, as is supposed for some species of *Caenorhabditis* [18]. However, we find no evidence of disproportionately lower within-male sperm size variability for species with larger sperm (phylogenetic contrasts on log-transformed values; F_1,23_=0.15, P=0.70) (Figure 5A), with the between-male CV and the phylogenetically un-corrected analysis actually showing a trend of higher CV in species with larger sperm (between-male PIC F_1,23_=4.47, P=0.045; Figure S4). These observations suggest that the net strength of selection on *Caenorhabditis* sperm size might be similar among species, with equally strong stabilizing selection favouring different optimal sperm sizes in different species.

**Figure 5.**
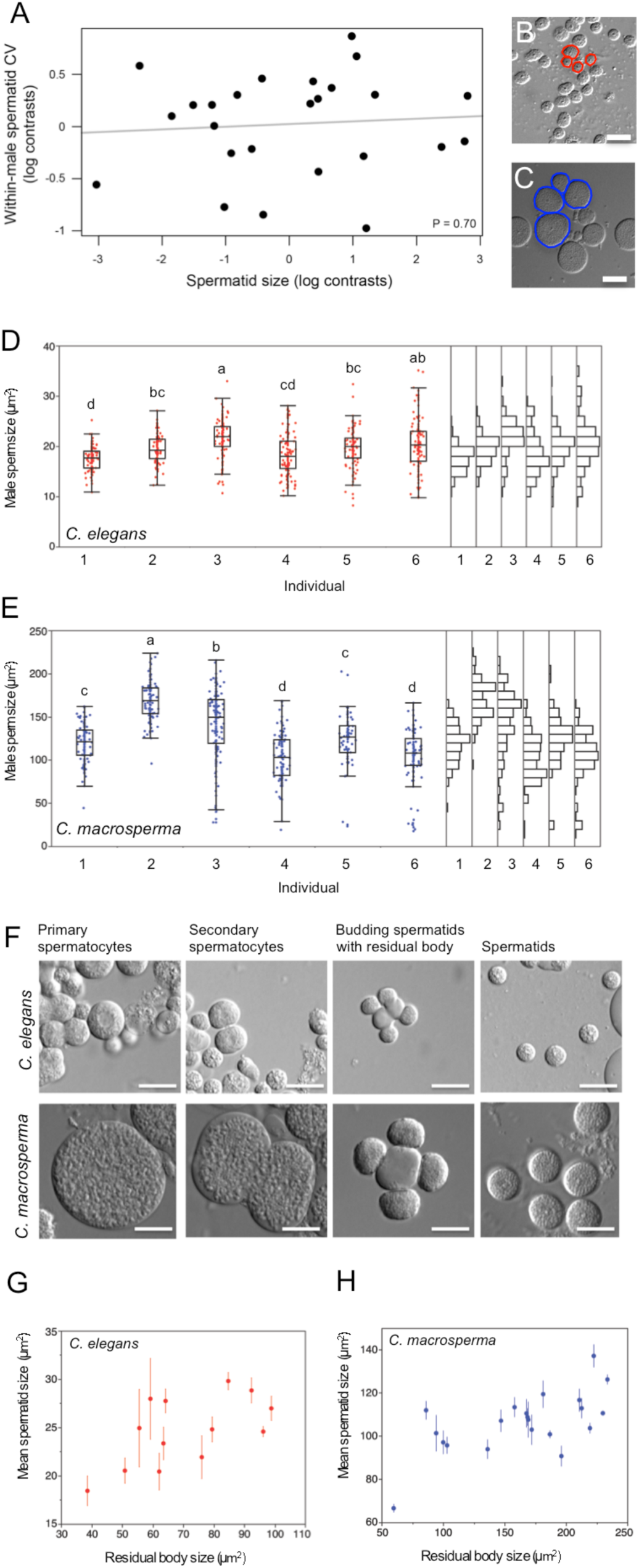
Intra-and inter-individual variation in male sperm size. (A) PIC plot of mean sperm size versus mean within-male CV of sperm size indicates no association (F_1,23_=0.15, P=0.70). (B,C) DIC images illustrating spermatid size variation within individual males of (B) *C. elegans* N2 and (C) *C. macrosperma* JU1857. Scale bars: 10μm. (D,E) Inter-and intra-individual variation in male sperm size in (D) *C. elegans* N2 (N=94-121 sperm per individual) and (E) *C. macrosperma* JU1857 (N= 74-138 sperm per individual). In both species, individuals show significant differences in average sperm size (*C. elegans* N2: F_5,647_=14.73, P<0.0001; *C. macrosperma* JU1857: F_5,646_=76.64, P<0.0001) and variance of sperm size (Levene’s Test, *C. elegans* N*2*: F_5,647_=9.29, P<0.0001; *C. macrosperma* JU1857: F_5,646_=10.43, P<0.0001). Values labelled with different letters indicate significant differences in mean sperm size (Tukey’s HSD, P<0.05). (F) DIC images of spermatocyte divisions in *C. elegans* N2 and *C. macrosperma* JU1857. Scale bars: 10μm. (G,H) Relationship between size of residual body and spermatid size (N=3-4/spermatocyte) originating from the same primary spermatocyte (mean ± sem) in (G) *C. elegans* N2 (F_1,11_=6.16, R^2^ =0.36, P=0.0305) and (H) *C. macrosperma* JU1857 (F_1,19_=12.95, R^2^ =0.41, P=0.0019).

A nested analysis of variance to partition variation among sources of male sperm size variation across the 26 *Caenorhabditis* species indicates that 5% of the total variation can be attributed to inter-individual differences, and 13% to intra-individual differences (Table S2). Moreover, comparison of intra-versus inter-individual variation in male sperm size suggests that intra-individual sperm size variation exceeds inter-individual variation in 42 of 47 isolates (21 of 26 species) analysed (Table S11). Two species with stark differences in average male sperm size, *C. elegans* (N2) versus *C. macrosperma* (JU1857), illustrate such pervasive sperm size variability within and between individuals irrespective of species mean sperm size (Figures 5B and 5C). In both species, differences among individuals of a single genotype are significant for both mean and variance of sperm size (Figures 5D and 5E). Intra-individual variation also is pronounced for *C. elegans* and *C. macrosperma*. For example, a single *C. macrosperma* individual may produce sperm that vary more than ten-fold in cross-sectional area, from approximately 20μm^2^ to over 200μm^2^ (mean 127.1±0.6μm^2^) (Figure 5E). As we observed for inter-and intraspecific variability in average sperm size, primary spermatocyte size shows similarly high variability within and between individuals (Figure 3C), consistent with the interpretation that sperm size variation originates primarily from size variation in primary spermatocytes (Figure 3D).

While variation in sperm size, whether due to genetic or environmental sources, appears to be explained predominantly by size variation in primary spermatocytes, additional intra-individual variation could potentially be introduced later in spermatogenesis. For example, size modification of spermatids derived from a single spermatocyte through asymmetric divisions during meiosis I and II could contribute to downstream heterogeneity in sperm size. To test this possibility, we examined meiotic division stages of male spermatogenesis in *C. elegans* N2 and *C. macrosperma* JU1857 (Figure 5F). Labelling with DAPI and antibodies for Actin and MSP (Major Sperm Protein) suggests evolutionarily conserved processes of sperm maturation, characterized by MSP-positive sperm cells and the exclusion of actin into the residual body [33, 42] (Figure S5). Quantifying size variation of spermatids derived from individual spermatocytes using DIC microscopy, we observed no evidence of consistent asymmetric meiotic divisions: spermatids budding off the same residual body are similar in size (Figures 5G and 5H). However, we cannot exclude that additional size variation of spermatids is occasionally introduced due to asymmetric resorption of residual body contents, as previously reported in *C. elegans* [31]. Nevertheless, residual body size correlates positively with average spermatid size (Figures 5G and 5H), lending further support to the conclusion that it is size variation of primary spermatocytes that provides the dominant source of size variation among spermatids (Figure 3D).

In *C. briggsae*, and perhaps other species, sperm bearing the X chromosome enjoy a fertilization advantage over nullo-X sperm [43, 44]. Differential sperm sizes owing to asymmetric meiotic cell division provides one possible mechanism for manifesting such an advantage, as occurs in a species of *Rhabditis* nematodes [45]. However, we detected no strong signal of bimodality in male sperm size distributions of individuals for *C. briggsae* or other species (Figures 5D, 5E and S6), arguing against an obvious size difference between X versus nullo-X sperm and being consistent with our previous results that sperm size determination occurs early, i.e. prior to the formation of haploid spermatids. Nevertheless, a more focused study is warranted to more thoroughly test the possibility of sex-chromosome dependent differences in *Caenorhabditis* acting on sperm size or other potential traits influencing sperm competition.

### Size changes during the transition from spermatid to spermatozoon

During the process of sperm activation, termed spermiogenesis, sperm cells undergo extensive morphological changes, marked by growth of a single pseudopod that is required for motility and which thus represents a presumed key element of sperm competitive ability [14, 46, 47]. We therefore asked whether spermatozoon morphology might differ between *Caenorhabditis* species with divergent spermatid sizes. All species examined displayed a spermatozoon phenotype similar to *C. elegans*, characterized by the polarization of the sperm cell into a cell body retaining the nucleus and other membranous cell components [27, 28] and the formation of a single pseudopod of variable shape (Figure 6A). The one exception was the spermatozoon morphology in *C. plicata* that, although clearly polarized, appear to retain a spherical shape without stereotyped pseudopod formation (Figure 6A); however, this atypical sperm morphology could be caused by increased sensitivity to the non-physiological conditions of the sperm medium, resulting in pseudopod retraction as observed for spermatozoa of other species after prolonged exposure to sperm medium (data not shown).

**Figure 6.**
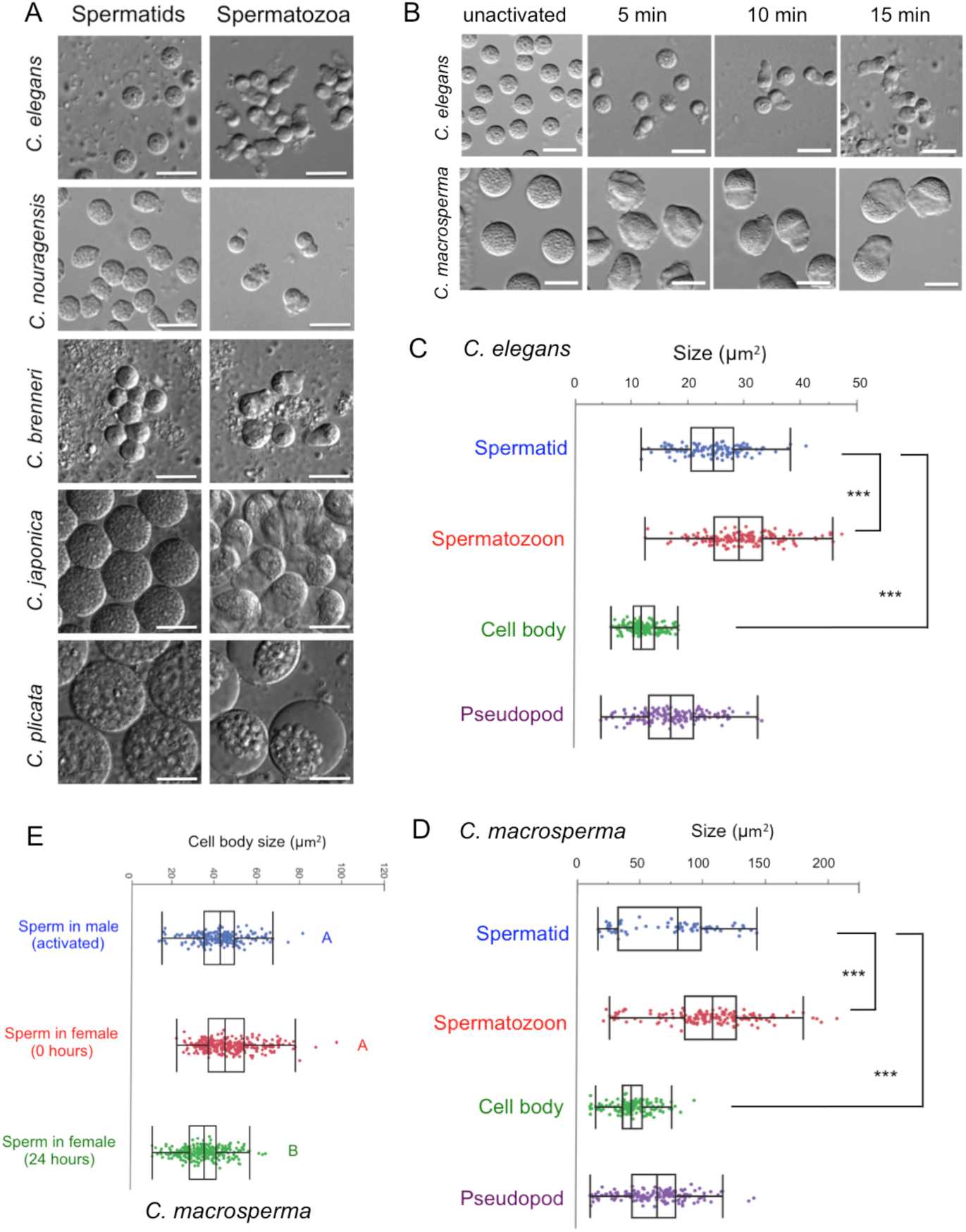
Size variation in spermatozoa. (A) DIC images of spermatids and spermatozoa (obtained from inseminated females) in *Caenorhabditis* species with divergent sperm size. Scale bars: 10μm. (B) DIC images taken during time course of Pronase in vitro sperm activation for *C. elegans* N2 and *C. macrosperma* JU1857. Scale bars: 10μm. (C,D) Size comparison of spermatids, spermatozoa (sum of cell body and pseudopod sizes), spermatozoon cell body, and pseudopod for *C. elegans* N2 and *C. macrosperma* JU1857 (D). Spermatozoon cross-sectional area is larger than for spermatids (ANOVA, *C. elegans* N2: F_1,263_=64.42, P<0.0001; *C. macrosperma* JU1857: F_1,208_=32.08, P<0.0001) and spermatozoon cell body size is strongly reduced relative to spermatid size (ANOVA, *C. elegans* N*2*: F_1,263_=654.48, P<0.0001; *C. macrosperma* JU1857: F_1,208_=73.84, P<0.0001). (E) Size distributions of spermatozoa (inferred from measurements of cell body size) in males (after in vitro activation) and in mated females (0h versus 24h after mating) in *C. macrosperma* JU1857. Average sperm size is significantly smaller in females 24h after mating compared to females immediately after mating (and activated sperm from males) (ANOVA, F_2,726_=88.83, P<0.0001). Values labelled with different letters indicate significant differences (Tukey’s HSD, P<0.05).

To test whether species differences in spermatid size affect spermatozoon morphology, we focused again on *C. elegans* N2 and *C. macrosperma* JU1857. In both species, in vitro sperm activation rapidly induces polarization of sperm cells, which progressively form a mature spermatozoon with a hemispherical cell body containing granular cell components and a single pseudopod (Figure 6B). Spermatid size differences between *C. elegans* N2 and *C. macrosperma* JU1857 translate into corresponding differences in size of cell body and pseudopod, and thus final size of spermatozoa (Figures 6C and 6D), suggesting that the entire size range of spermatids activate successfully into mature spermatozoa. Moreover, we observed no difference between the species in the size ratio of pseudopod to cell body (ANOVA, F_1,298_=0.23, P=0.63), suggesting that relative sizing of cell body versus pseudopod formation is maintained irrespective of spermatid size. These measurements, although limited to analysis with DIC microscopy, further suggest that the cell body size of spermatozoa becomes substantially reduced, by a factor of two on average, relative to the spermatid size (Figures 6C and 6D).

The above data indicate that *Caenorhabditis* females inseminated by a single or multiple males will contain a pool of available spermatozoa that varies substantially in size. Under size-dependent sperm competition, the sperm size distribution in a female’s reproductive tract should thus shift over time as larger sperm take precedence in fertilization and leave smaller remaining sperm at later time points. Consistent with this prediction, we found in *C. macrosperma* that females retain significantly smaller complements of sperm 24 hours after mating compared to the sperm size distribution found in their reproductive tracts immediately following mating or derived directly from males (ANOVA, F_2,726_=88.83, P<0.0001) (Figure 6E).

## DISCUSSION

### Convergent evolution of sperm size gigantism in *Caenorhabditis* nematodes

Our phylogenetic analyses of sperm size evolution, including many novel *Caenorhabditis* species, uncover surprisingly high disparity in male sperm size among gonochoristic species that reveal multiple independent origins of sperm gigantism. The recurrent evolution of exceptionally large sperm has been reported for diverse taxonomic groups, including invertebrates (e.g. insects and molluscs) and vertebrates (e.g. mammals and birds) [5-8, 48]; however, in contrast to *Caenorhabditis* nematodes, most of these taxa possess flagellate sperm. While our observations are consistent with the idea that *Caenorhabditis* sperm size divergence reflects the product of sexual selection by sperm competition, it will be critical to determine how characters of the female reproductive morphology associate with observed sperm size variation to evaluate the significance of coevolutionary processes between sperm size and female reproductive tract [26].

Moreover, how specific ecological or life history factors of different *Caenorhabditis* species might have shaped such differential investment for many small versus fewer large sperm remains unresolved, given our limited but expanding knowledge of *Caenorhabditis* natural history [19, 49, 50]. Notably, however, two of the four species with gigantic sperm display specialized life histories: *C. japonica* and *C. drosophilae* have dispersal associations with specific phoretic host insects [51-53]. The two other species with gigantic sperm (*C. macrosperma* and *C. plicata*) also potentially represent “specialists” given that *C. macrosperma* displays a very localized geographic distribution [54] and that *C. plicata* was isolated only once and is the only *Caenorhabditis* in lab culture that was isolated from carrion [52]. This limited evidence suggests that specialist life histories might be predisposed to conditions favourable to the evolution of extremely large male sperm. The life cycle in nature of *C. japonica* is known in most detail, for which reproduction takes place in isolated populations among the tens of founding individuals [53]. Because male reproductive success will be determined by the product of the number of mates and the number of fertilizations per mate, the relative importance of sperm size versus number in male fitness will depend on whether mate number or fertilization success per mate most constrains male fitness. We hypothesize that, in such tiny mating groups, male reproductive success might be constrained less by number of mates than by successful fertilization of their multiply-inseminated mates, thus putting a fitness premium on the competitiveness of individual sperm and favouring the evolution of exaggerated sperm size. In contrast, we hypothesize that for most *Caenorhabditis*, a larger number of mates inseminated provides a greater relative benefit to male fitness and thus favours the evolution of especially vigorous mating ability and rapidly-produced, numerous sperm at the expense of sperm being individually modest in size.

### Evolution of reduced sperm size in androdioecious species

Male sperm size evolution in androdioecious species provides conditions of especially weak selection on male-male sperm competition, owing to male rarity in populations [49, 55, 56]. Thus, in addition to relaxed selection on male reproductive performance in general [57, 58], the convergent evolution of miniature sperm in androdioecious species likely results from a combination of: (i) selection favouring small sperm under low sperm-competition risk conditions (both sexes), (ii) selection for enhanced hermaphrodite self-fertilizing reproductive success that disfavours resource allocation to sperm (hermaphrodites only), (iii) developmental biases of somatic-sex causing sperm size reduction (hermaphrodites only) [17]. Interestingly, our finding of limited evidence for correlated sperm size between hermaphrodites and males of a given species suggests a genetic decoupling of sperm size determination between the sexes in this sexually-dimorphic trait. Despite the special selective forces on male reproductive traits in androdioecious species, we also found that males of some gonochoristic *Caenorhabditis* have similarly miniature sperm (*C. nouraguensis, C. yunquensis, C. angaria, C*. sp. 1, *C. castelli*). As a consequence, future research on sperm size evolution in this group should aim to understand the factors that drive sperm miniaturization as well as gigantism. It remains unknown whether these factors might be more likely to involve mating-group size dynamics that maximize sperm count rather than size or an increased role of seminal fluid components in sperm competition, as analysis of *comp-1* mutants in *C. elegans* has made it clear that sperm size is not the sole determinant of sperm competitive ability [47].

### Consistently high levels of intra-and inter-individual sperm size variability

For all *Caenorhabditis* species examined, we find previously underappreciated high levels of intraspecific sperm size variability that includes pronounced inter-and intra-individual sperm size heterogeneity. Significant variation in sperm morphometric traits between and within individuals is also common in diverse taxa that produce flagellate sperm [59, 60]. However, in contrast to our observations for *Caenorhabditis*, within-individual sperm size variances in these other taxa were generally low and smaller than size variance calculated among different individuals [59-62]. In general, the evolutionary significance and developmental origins of inter-and intra-individual sperm trait variance are largely unknown, perhaps with the exception of some insect species that display pronounced sperm heteromorphism associated with functional differentiation of distinct sperm forms [63-65]. While our data provide no obvious signature of discrete size classes of sperm with different functions, it remains to be determined whether sperm across the entire size range of an individual are all fully functional, and whether the behavior of an individual sperm depends on its size. The presence of consistently high intra-individual sperm size variance across all *Caenorhabditis* species and genotypes – largely irrespective of mean sperm size – suggests several potential underlying causes. For example, (i) the maintenance of low sperm size variance may be costly, e.g. because increased precision would come at the cost of reduced sperm production speed [38, 39], (ii) increased sperm size variance reflects an adaptive strategy to maximize both mean and number of sperm produced, or (iii) the developmental architecture of spermatogenesis constrains the precision with which sperm trait size can be achieved. Experimentally evaluating the relative contributions of such adaptive and non-adaptive forces in the determination of sperm size variability should thus be a key priority for future research.

### Evolution of sperm trait correlations, trade-offs, and significance of sperm gigantism

In this study, we have focused on size in sperm evolution. However, male allocation of ejaculate expenditure per mating comprises not only sperm size, but also the number of sperm and the amount and composition of the non-sperm seminal fluid [66]. In *Caenorhabditis*, sperm size and number trade off such that genotypes that produce larger sperm also make them at a slower rate [14, 23] and, as we show here, transfer fewer per mating (Figure 2F). Despite the greater competitive ability of larger sperm within a reproductive tract, it remains unresolved what, mechanistically, is most critical for securing the competitive advantage: is it the greater speed conferred by a larger pseudopod [14, 46, 47], or might it be better adhesion capability to the interior walls of the uterus and spermathecae, or a greater capacity to dislodge smaller sperm from the best locations? Resolving these possibilities would shed more light on the details of sperm competition traits most subject to selection, and the molecules underpinning them.

*Caenorhabditis* males, however, also transfer a largely unknown mixture of seminal fluid components and deposit a copulatory plug upon mating. The plug is comprised primarily of the mucin protein PLG-1 with possible functions including mate guarding and sperm retention [47, 67-70]. It remains to be discovered how these non-sperm components of the ejaculate affect fertilization success and whether they might also contribute a source of tradeoffs in reproductive resource allocation (e.g. plug size, seminal fluid quantity or complexity) for overall ejaculate expenditure.

A consequence of sperm gigantism in *Caenorhabditis*, in the absence of correspondingly large oocytes, is the drastic reduction in the magnitude of anisogamy. We estimate that 3-5% of the volume of the zygote will derive from the sperm in such species, in contrast to <0.5% for the sperm contribution to the zygote for species at the other extreme that have minute sperm. How efficient will the maternally-provisioned proteasome and RNA degradation machinery be in the face of such a large influx of cytoplasmic material, given that dogma holds that sperm contribute only the haploid complement of chromosomes and centrioles to the zygote? Might changes in sperm-mediated anisogamy resulting from male-male sperm competition produce an arena for novel selective pressures? Two hypotheses immediately spring from the consequences of a large cytoplasmic input from extremely large sperm. First, could the sperm’s cytoplasm provide a resource, provisioning ‘nutrients’ to the developing zygote? For example, a variety of insect males deliver seminal ‘nuptial gifts’ to their mates that can act as a food or water resource that fosters female reproduction [8, 71], so it is conceivable that exceptionally large sperm might represent a more direct route for a paternal energetic contribution to their offspring. Second, might the sperm pack a suite of molecular and cellular components that engage in parental sexual conflict over the control of gene regulation in early zygotic development? In *Caenorhabditis*, such sexual conflict might be mediated by maternal and paternal contributions of small RNAs [72] or protein products, with paternal-effect examples including PEEL-1 [73]. The significance of these mechanisms in mediating potential sexual conflict, and whether they can be linked to differences in sperm size, remains to be addressed. Moreover, because *Caenorhabditis* offspring production usually is not viviparous, models of parental conflict from mammals and plants largely do not apply [9]. However, sexual antagonism in the control over sexual development in the embryo provides an arena for paternal interests to manifest the inter-sexual conflict [9, 74].

### Developmental origins of *Caenorhabditis* sperm size variation

The developmental mechanisms regulating sperm size, in both flagellate and aflagellate taxa, are generally poorly understood. Our study shows that *Caenorhabditis* sperm size is largely determined during early spermatogenesis, and specifically implicate heterogeneity in the size of primary spermatocytes as establishing the developmental basis of variation in sperm size. By contrast, subsequent cell divisions introduce little sperm size variability, as would result if the birth of spermatids during meiosis II cell divisions were asymmetric. The key role of primary spermatocyte size in controlling subsequent sperm size holds true for comparisons between species, between genotypes within species, and even within individuals and between the sexes for those androdioecious species in which both males and hermaphrodites produce sperm. These results suggest that *Caenorhabditis* sperm size is established at the diploid stage, similar to previous studies, which did not find any evidence for haploid determination of sperm length in flies [75].

Why does *Caenorhabditis* sperm size determination occur so early during development and what is it that sets the size of primary spermatocytes? The beginning of the growth phase of spermatocytes coincides with the transition from pachytene into diplotene stages of meiosis [30] (Figure 3B). Growth of spermatocytes seems to occur throughout the progression through this condensation zone until entry into metaphase when spermatocytes detach from the rachis, the gonad core with shared cytoplasm, and start to divide [30]. The beginning of the growth phase occurs roughly at the same stage (diakinesis) in *C. elegans* oocytes although meiotic progression of sperm cells is considerably (2-3x) faster than for oocytes [76], consistent with gamete size differences. Further mechanisms of growth control for spermatocytes and oocytes are distinct, at least in part: streaming of cytoplasmic material from the gonad syncytium into growing oocytes, i.e. a likely provisioning mechanism, does not occur during spermatogenesis [77]. In addition, ‘physiological’ germ cell apoptosis – thought to generate resources for developing oocytes – does not occur during spermatogenesis of *C. elegans* males or hermaphrodites [78]. Overall, then, early size determination of *Caenorhabditis* can be explained by the limited growth phase corresponding to the time of primary spermatocyte formation and when spermatocytes are still connected to the gonad rachis, potentially providing nutrients. After detachment from the rachis, spermatocyte growth may thus become impossible, and/or meiotic divisions occur too rapidly to allow for significant cell growth.

Given the extreme size disparity of spermatocytes and oocytes in *C. elegans*, spermatocyte growth may not require oocyte-like mechanisms of gamete provisioning. However, this situation may not hold for those *Caenorhabditis* species that produce much larger sperm. It remains to be tested whether spermatogenesis in large-sperm species might have co-opted oocyte-like mechanisms to nurture sperm growth. Fundamental mechanisms regulating cell size, and thus also spermatocyte size, include growth rate and timing of cell cycle progression [79]; for example, if spermatocyte growth rate remains constant, simply delaying entry into metaphase may lead to increased cell size. Consistent with this scenario, sperm production rates are slower for *C. elegans* genotypes that make larger sperm [14, 23]. Further experimental analysis may permit the disentangling of the interplay among the diversity of potential extracellular factors influencing spermatocyte growth with cell-autonomous mechanisms involved in *Caenorhabditis* sperm size regulation.

## Experimental Procedures

### Nematode strains and cultivation

Strains were maintained at 20°C on 2.5% agar NGM (Nematode Growth Medium) plates seeded with the *E. coli* strain OP50 [80]. The following species (strains) were used in this study: *C. angaria* (PS1010), *C. brenneri* (CB5161, JU1398, SB280), *C. briggsae* (AF16, ED3092, HK104, JU1341, QR24), *C. drosophilae* (DF5077), C. *elegans* (CB4856, JU258, LKC34, MY2, N2), *C. japonica* (DF5081), *C. plicata* (SB355), *C. remanei* (PB4641, SB146, VT733), *C*. sp. 1 (SB341), *C. doughertyi* (JU1333), *C. tropicalis* (JU1373, JU1630, JU1818, NIC58, QG131), *C. castelli* (JU1427), *C. virilis* (JU1528), *C. imperialis* (EG5716, NIC118), *C. kamaaina* (QG122), *C. wallacei* (JU1873), *C. nouraguensis* (JU1825), *C. macrosperma* (NIC293, NIC401, JU1857,) *C. yunquensis* (EG6142), *C*. sp. 2 (DF5070), *C. guadeloupensis* (NIC113), *C. sinica* (JU800), *C. portoensis* (EG4788), *C. afra* (JU1199), *C*. sp. 8 (DF5106, NIC184, QX1182) and *C. nigoni* (EG5268). *C. macrosperma* isolates NIC293 and NIC401 were isolated in French Guiana in 2013 (CB and ADC, unpublished data). For detailed strain information, see references [34, 52, 54].

### Sperm size measurements

Males were isolated from strain cultures at the L4 stage and maintained on NGM plates containing males only, to obtain spermatid size measurements from synchronized and unmated males. After 24 hours at 20°C, when males had reached the adult stage, spermatids were obtained by needle dissection of males in sperm medium (50mM HEPES pH7.8, 50mM NaCl, 25mM KCl, 5mM CaCl_2_, 1mM MgSO_4_, 1mg/ml BSA) [81]. Spermatids from multiple males were immediately imaged using DIC microscopy (60x or 63x objectives). Using ImageJ software (Rasband, W.S., ImageJ, U. S. National Institutes of Health, Bethesda, Maryland, USA, http://imagej.nih.gov/ij/, 1997-2014), we calculated length and width of each spermatid to obtain measures of cross-sectional area assuming an ellipse shape: π x (length/2) x (width/2) (measured at approximately 1000x magnification). Hermaphrodite spermatids were dissected from young unmated adults (mid L4+24h), and imaging and spermatid size calculations were performed as described above. At this developmental stage, most hermaphrodite individuals contained both spermatids and activated sperm (spermatozoa), and the latter were thus not included for measurements.

### Body size measurements

Estimates of body length and width were obtained by measuring adult males and females (hermaphrodites) at L4 + 24h. Live individuals were on 4% agarose pads in M9 buffer, containing 100mM sodium azide [80] and imaged using a 10x objective. Using ImageJ software, we measured the body midline (from mouth to tip of tail) to estimate length and we measured body width in the mid-section of adult animals perpendicular to the anterior-posterior axis.

### Anisogamy measurements

To estimate egg size we used embryo size measurements obtained by Farhadifar et al. (2015), with cross-sectional area calculated assuming an ellipse from length and width values. The index of anisogamy was calculated as egg volume divided by spermatid volume, where egg and spermatid volumes presume an ellipsoid cell shape: (4/3) x π x (length/2) x (width/2)^2^.

### Mating experiments and sperm number measurements

To quantify numbers of transferred sperm after a single mating for three independent contrasts of species pairs differing in sperm size, we followed the phylogeny established by Kiontke et al. (2011) (Figure 2F). For each strain, males and females were picked at the L4 stage and maintained on separate NGM plates for 36h at 20°C. A single virgin female and five unmated males were then placed together on individual mating plates (*E. coli* OP50 lawn of 5 mm diameter) and observed at 80x magnification using a dissecting microscope. As soon as a single mating event had been completed, i.e. spicule insertion and ejaculation (visualized as sperm flow from the male *vas deferens* into the female uterus) and after which the male had left the female, the inseminated female was isolated and fixed in ice-cold Methanol. Females were then washed twice in M9 buffer and mounted in DAPI-containing Vectashield medium (Vector Laboratories, Inc., Burlingame, CA, USA). Imaging and sperm counts were performed as previously described [82]. In brief, images were taken at 40x magnification as Z-stacks covering the entire thickness of the animal using an epifluorescence microscope. We then manually counted sperm number (in uterus, spermatheca, proximal germline) by identifying condensed sperm nuclei of each focal plane using the ImageJ plugin Cell Counter (Rasband, W.S., ImageJ, U. S. National Institutes of Health, Bethesda, Maryland, USA, http://imagej.nih.gov/ij/, 1997-2014).

### Primary spermatocyte measurements

Because individual primary spermatocytes are difficult to isolate and not easily staged using DIC microscopy, we estimated cell size of undissected male primary spermatocytes in the karyosome stage at the end of meiotic prophase, when DNA content is highly condensed [30], using DAPI to stain nuclei and Phalloidin to visualize cell outlines (Figure 3B). Males were isolated from strain cultures at the L4 stage and maintained on NGM plates containing males only, to obtain spermatid size measurements from synchronized and unmated males. After 24 hours at 20°C, male gonads were extruded in M9 buffer supplemented with levamisole using syringe needles. Extruded gonads were fixed in 4% paraformaldehyde for 10 minutes followed by a 5 min PBS-Triton X-100 (0.1%) wash at room temperature. Extruded gonads were stained for actin using Phalloidin (1:500 dilution, Sigma-Aldrich) overnight at 4°C in a humidified chamber. Gonads were washed in PBS and mounted in Vectashield mounting medium supplemented with DAPI. Images of the germline section containing primary spermatocytes were obtained using an epifluorescence microscope (40x objective). Size measurements were restricted to cells where the polygonal cell outline was completely visible and where DNA was highly condensed. Measurements of spermatocyte area were obtained by delineating the circumference of cells using ImageJ software.

Measurements of primary spermatocyte size were obtained from the following strains: *C. elegans*: N2, *C. guadeloupensis*: NIC113, *C. remanei*: PB4641, *C. yunquensis*: EG6142, *C. brenneri*: CB5161, *C. japonica*: DF5081, *C. drosophilae*: DF5077, *C. plicata*: SB355 and *C. macrosperma*: JU1857.

### Sperm activation assays and measurements of spermatozoa

In vitro sperm activation assays in *C. elegans* N2 and *C. macrosperma* JU1857 (Figures 6B to 6D) were performed by dissecting male spermatids in sperm medium supplemented with Pronase E (Sigma-Aldrich), as previously described [83, 84]. Images of spermatozoa were taken within15-20 min after dissection, using DIC microscopy (60x or 63x objective). Size estimates (area) were obtained my measuring the circumferences of cell body and pseudopod of each spermatozoon using ImageJ software; total spermatozoon size was calculated as the sum of cell body plus pseudopod.

### Measurements of C. macrosperma spermatozoa after mating

To test for sperm size differences between males, females (0h after mating) and females (24h after mating) (Figure 6E), we isolated *C. macrosperma* JU1857 males and females at the L4 stage, and kept them on separate plates to prevent mating. After 24 hours, 10 males and 5 virgin females were transferred to each of several mating plates (*E. coli* OP50 lawn of 5 mm diameter). After 5 hours, both males and females were dissected in sperm medium to obtain spermatids and spermatozoa, respectively. Spermatids dissected from males were activated using Pronase and measured after 15-20 minutes. Sizes of spermatid and spermatozoon (cell body and pseudopod) were measured as described above. Additional females from the same experiment were maintained without males for another 24 hours prior to dissection of spermatozoa. Size measurements were done in the same fashion as outlined above.

### Phylogenetic analysis

We performed phylogenetic independent contrasts (PIC) on species mean trait values (log-transformed) for the *Caenorhabditis* phylogeny and branch lengths from [34], as implemented in the R package APE [85]. Body size (male and female length and width) and egg (embryo cross-sectional area) measurements used species mean values from measurements described above. For sperm size, we used least-squares mean estimates of spermatid cross-sectional area for each species from a generalized linear model that incorporated variation within species in our spermatid size measurements owing to strain and individual. PIC analyses were performed on log-transformed trait values to eliminate scale-dependence. We also applied SURFACE [36] to test for the number of independent and convergent shifts in sperm size and embryo size along the phylogeny, which uses the Akaike Information Criterion (AIC) for model selection of the number of phenotypic regimes under Ornstein-Uhlenbeck process using a Hansen model of trait evolution along the phylogeny. To complement AIC, we also conducted 500 phylogenetic trait simulations in SURFACE to derive a P-value for the inferred number of regime transitions (*c*) for sperm size. For graphical mapping of ancestral state inference, however, we applied GEIGER [86], which uses a Brownian motion model of trait change on the phylogeny. Metrics of phylogenetic signal (Pagel’s λ, Blomberg’s k) were calculated in R using the GEIGER and PICANTE packages [87]. All R scripts are publicly available at http://github.com/cutterlab.

## Acknowledgements

*Caenorhabditis* wild isolates were kindly provided by Karin Kiontke, Marie-Anne Félix, Michael Ailion, Matt Rockman, and the *Caenorhabditis* Genetics Center. CB acknowledges financial support by the Centre National de la Recherche Scientifique (CNRS), the Agence Nationale de la Recherche (ANR) and the Fondation Schlumberger pour l’Education et la Recherche (FSER). ADC received support from the Natural Sciences and Engineering Research Council (NSERC) of Canada and from a Canada Research Chair. NC was supported by a fellowship of the France Canada Research Fund (FFCR).

## Author contributions

Conceived and designed the experiments: CB and ADC. Performed the experiments: AV, NC, CG, NP. Analysed the data: CB, ADC, AV, and JCG. Wrote the paper: CB and ADC.

## Competing Interests

The authors declare no competing interests.

